# Aging reduces excitatory bandwidth, alters spectral tuning curve diversity, and reduces sideband inhibition in L2/3 of primary auditory cortex

**DOI:** 10.1101/2025.04.02.646797

**Authors:** Kate Maximov, Patrick O. Kanold

**Author notes:** **Corresponding authors:** Patrick O. Kanold, Dept. of Biomedical Engineering, Johns Hopkins University, Miller 380. Baltimore, MD 21205 USA.

## Abstract

Presbycusis, or age-related hearing loss, is caused by changes in both the peripheral and the central auditory system. Many of the peripheral structures that degrade with age have been identified and characterized, but there is still a dearth of information pertaining to what changes occur in the aging central auditory pathway that are independent of peripheral degradation. The primary auditory cortex (A1) of aging mice shows reduced suppressive responses and reduced diversity of temporal responses suggesting alteration of inhibitory processing. To gain a better understanding of how tuning features of the auditory cortex change with age, we performed *in vivo* 2-photon Ca^2+^ imaging on L2/3 of the auditory cortex of both adult (n=14, 11-24 weeks old) and aging (n=12, 12-17 months old) mice that retain peripheral hearing in old age. To reveal inhibitory inputs to L2/3 neurons we characterized spectral receptive fields with pure tones and two tone complexes. We find that in contrast to adult mice, L2/3 excitatory neurons from aging mice showed fewer distinct categories of spectral receptive fields, though in a subset of FRA types, we found increased diversity. We also noted a decrease in excitatory bandwidth with age among broadly tuned neurons, but that sideband inhibition became weaker across all FRA types due to a reduced amplitude in inhibitory responses. These results suggest that aging causes changes in circuit organization leading to more homogenous spectrotemporal receptive fields and that the lack of response diversity contributes to a decreased encoding capacity observed in aging A1.

## INTRODUCTION

It is estimated that over 50% of adults over 60 years old have meaningful hearing loss (Lin, Thorpe et al. 2011). Hearing loss can have widespread health ramifications, extending beyond communication troubles to withdrawal from society and peer interactions and is comorbid with depression, dementia, and stroke (Deal, Reed et al. 2019, Huang, Jiang et al. 2023). Structures in both the peripheral and central levels of the nervous system degrade with age and contribute to age-related hearing loss (Gopinath, Rochtchina et al. 2009, Lin, Thorpe et al. 2011), but several studies have shown that deficits can happen at the central level, independent of noticeable changes in the periphery, as adults with hearing difficulty can present with functionally intact peripheries (Fitzgibbons and Gordon-Salant 1998, Gordon-Salant and Fitzgibbons 1999).

Studies of aging mice have shown that excitatory neurons in primary auditory cortex (A1) have less suppressed responses, show less sensitivity to temporal changes, and have increased correlated activity (Shilling-Scrivo, Mittelstadt et al. 2021). Moreover, it has been shown that receptive fields in A1 become more complex with age and neurons fire less reliably (Turner, Hughes et al. 2005). Such changes on the neural level could be caused by a reduction of inhibition in the aging A1. Indeed, multiple studies have suggested that aging is associated with reduced inhibition. Studies in rodents have shown that expression of inhibitory markers e.g. parvalbumin is reduced in the aging brain (Stanley, Fadel et al. 2012, Ouellet and de Villers-Sidani 2014, Brewton, Kokash et al. 2016). Thus, we hypothesized that such changes to inhibition should be present at the functional level. Neurons in A1 can show complex spectral receptive fields which are shaped by the interplay of excitatory and inhibitory inputs (Tan, Zhang et al. 2004, Li, Ji et al. 2014, Zhou, Liang et al. 2014, Meng, Winkowski et al. 2017, Liu and Kanold 2021). While inhibitory inputs are not easily resolved when using pure tone stimuli due to the low baseline firing rates of A1 neurons, two tone stimuli can be used to reveal inhibitory inputs. In this study, we determined to characterize the age-related functional changes that occur in the excitatory neurons of the central auditory system of mice. By using mice on a CBA/CaJ background, we were able to isolate our observations to central changes only, as this strain retains peripheral hearing in old age (Willott, Parham et al. 1988, Spongr, Flood et al. 1997, Bowen, Winkowski et al. 2020, Shilling-Scrivo, Mittelstadt et al. 2021, Park, Willott et al. 2024).

We used in vivo 2-photon imaging with pure tone and two tone stimuli to measure responses of A1 neurons and reveal inhibitory sidebands in both aging and adult animals. We find that neurons from aging animals show reduced functional diversity in their spectral responses, decreased bandwidth, and that inhibitory sidebands weakened. These results indicate that aging alters both excitatory and inhibitory processing in A1.

## METHODS

### Animal procedures

All procedures were approved by the Johns Hopkins University Animal Care and Use Committee. We used 12 aging mice (7 males, 5 females; age, 12-17 months) in our present study and 14 adult mice (8 males, 6 females; age, 11–24 weeks). We used a cross between CBA/CaJ mice (stock #000654, The Jackson Laboratory), which are known for retaining good hearing into old age (Willott, Parham et al. 1988), and Thy1-GcaMP6s mice (stock #024275, The Jackson Laboratory) which are used for calcium imaging of excitatory neurons, to create Thy1-GCaMP6s X CBA mice. All mice were tested for the early hearing loss mutation Cdh23^ahl^, which is the recessive allele that causes peripheral hearing loss and only mice with at least one normal copy were used in the study.

### Cranial window implant

We implanted cranial windows over left A1 to be able to perform calcium imaging following procedures previously outlined (Liu, Whiteway et al. 2019, Liu and Kanold 2021). 2 hours before surgery, mice were injected with dexamethasone (5 mg/kg) which prevents brain swelling and inflammation. The animals were anesthetized with isoflurane (Fluriso, VetOne) using a calibrated vaporizer (Matrx VIP 3000, Midmark) with 4-5% for induction and 1.5–2% during the surgical procedure. During surgery, the body temperature of the animal was maintained between 35.0 and 36.0°C. The head was fixed by placing the front teeth in a headfix bar and securing the nose. Next, the hair was removed first using scissors and then by applying Hair Remover Face Cream (Nair). After hair removal, a series of 3 applications of betadine then 70% ethanol was applied to sterilize the skin. The skin was then removed to expose the top and left sides of the skull – just anterior to bregma and slightly medial to the mandibular condyle. The surface of the skull was covered in hydrogen peroxide to dissolve soft tissue and was then scraped with a scalpel to remove remaining tissue and increase surface area for later cement application. Muscles covering the left temporal bone were subsequently retracted ∼3 mm from the ridge of the left parietal bone and any amount not able to be easily retracted was removed. After cleaning the skull, a custom 3D printed stainless steel headplate was mounted and secured using C&B-Metabond (Parkell). A circular craniotomy was then performed over the left auditory cortex with a diameter of 3.5 mm using a dental drill. Viral injections were performed at this point if necessary. Then a custom-made cranial window was placed over the exposed brain. The window consisted of two layers of 3 mm round coverslips (catalog #64–0720-CS-3R, Warner Instruments) stacked at the center of a 4 mm round coverslip (catalog #64–0724-CS-4R, Warner Instruments) and secured with optic glue (catalog #NOA71, Norland Products). The edge of the cranial window was then sealed with Kwik-sil (World Precision Instruments). More Metabond was then applied to secure the window to the skull. After the surgery, 0.05 ml Cefazolin (1 g/vial, West Ward Pharmaceuticals) was injected subcutaneously, and the animal recovered under a heat lamp for 30 min before being returned to the home cage.

### Widefield Imaging

Widefield imaging was conducted to locate areas responsive to sound within the cranial window. Mice were headfixed under an sCMOS pco.edge camera with a Nikon 4x objective and the window was illuminated with a blue LED (470 nm). A series of 16 tones ranging from 4 to 64 KhZ was presented 5 times each and fluorescent changes due to sound presentation were recorded. We used custom MATLAB software (Liu, Whiteway et al. 2019) to identify sound responsive regions and used the known tonotopic organization to identify auditory areas (Kanold, Nelken et al. 2014).

### 2 photon imaging

Imaging was performed as previously described (Liu, Whiteway et al. 2019). The tonotopic map collected from widefield imaging was used to find primary auditory cortex (A1) using 2 photon Ca2+ imaging. After locating A1, the z location was zeroed at the brain surface, and then an FOV was chosen between 170 and 210 um to be within cortical layer 2/3. Images were 512×512 pixels at 2x magnification and captured at 30 Hz.

### Sound stimuli

We presented 16 logarithmically spaced tones between 4 and 64 kHz at 3 different decibel levels (40, 55, 70) to obtain tuning curves from A1 neurons. Since excitatory neurons in A1 have low baseline firing rates, we additionally presented two tone combinations (i.e. 4&4.8kHz, 4&5.7kHz, etc) to characterize sideband inhibition by noting changes in the response level to that of the neuron’s pure tone response which could be attributed to a facilitative or inhibitory effect of a second tone. All single tones were calibrated at 70 dB and all tones used in two tone combinations were calibrated to 60 dB so the resulting sound played was 63 dB SPL. Each tone was 100 ms with a 10 ms linear ramp. All stimuli were generated using custom MATLAB code (Liu and Kanold 2021) that utilized a TDT RX6 and PA5 for waveform generation. Stimuli were played through a TDT ES1 speaker positioned roughly 10 cm from the animal’s right ear.

### 2 photon analysis

We detected neurons and extracted neural fluorescence traces using suite2p. We calculated neuropil corrected traces with the following equation F_corrected_ = F_cell_ – 0.8 * F_neuropil._ After neuropil correction, the full imaging session was separated into trials. We then calculated ΔF/F by subtracting the average value of 10 baseline (pre stimulus) frames from each post stimulus frame and then dividing the stimulus frames by the baseline as such ΔF/F = (F_corrected_ (t) - baseline) / baseline. Each “trial” began 10 frames before the sound onset frame and ended 35 frames after sound onset. Response significance was determined using prior methods (Liu, Whiteway et al. 2019, Liu and Kanold 2021). In brief, a window of 10 frames before stimulus onset was compared to a 20 frame window (spanning frames 10 to 30) after stimulus onset. A response was deemed significant if there was no overlap between the 99.9% confidence intervals across trials. After finding all significant responses, a frequency response area (FRA) could be constructed for each neuron, representing the neuron’s receptive field in frequency and decibel space.

### Kmeans classification and t-SNE visualization

We used an unsupervised classification algorithm to understand the diversity and recurring shapes of FRAs across groups. We chose to use an unsupervised algorithm over manual clustering so as not to bias results in comparing our adult and aging groups. Since FRAs can vary widely in shape and location in stimulus space (Liu and Kanold 2021), FRAs were aligned at their geometric center so that response shapes could be compared without regard to location in frequency space. For each neuron, nonsignificant responses were set to zero to obtain clearer shapes from clustering results. We then reshaped each neuron’s 3×16 (dB x frequency) matrix to 1×48 and added zero padding to each side of the vector so our ultimate matrix contained one vector for each neuron with all geometric centers aligned at the center of the larger matrix. We applied PCA to the dataset and kept the principal components that explained 95% of the variance. Next, we applied kmeans clustering to the reduced data and chose 6 clusters based on a within cluster sum of squares elbow plot. Finally, we used a t-SNE plot to visualize the clustered data. T-SNE plots are frequently used to complement PCA as they can reveal nonlinear structures that PCA can overlook and may show clusters that would not appear when plotting clustered data in PCA space. However, it is important to note that t-SNE plots focus on local relationships and can distort global patterns. Thus, there is no meaning in the location of clusters near each other, only to the density and location of points within each cluster which shows the datapoints’ similarity.

To find the compactness of each cluster, we calculated the Euclidian distance between each point and the center of its respective cluster. We then averaged over all values to find the “compactness score” – a larger score meaning on average greater distances between any neuron to its cluster center and hence a larger variability of points within the cluster.

### Receptive field variability analysis

We quantified and visualized the extent of the variability in responses in frequency or sound level space. We sought to compare changes over both frequency and sound level across the entire receptive field between groups. This was to ensure the inclusion of multipeaked neurons which can have significant responses outside of the main receptive field (Sutter and Schreiner 1991). To compare across different neurons within the same cluster shape, we first took the maximum value of each trial averaged responses for each SPL/frequency combination (Fig 4A, left and middle panel). We binarized the responses to 1 if the value was more than 30% of the maximum response of the whole FRA or 0 if the response fell below 30% (Fig 4A, right panel). We then summed these responses over frequency or SPL for all neurons and then computed the z-score. We then plotted the cumulative summed z-score of the responses across either frequency or SPL space (Fig 4B,C,D,E). The cumulative sum across a neuron’s receptive field gives us an idea of its response profile as the area across which significant responses occur dictates the slope of the curve. More sparsely tuned types will have a steeper slope as there are fewer frequencies contributing to the shape and all are localized at the center of the FRA, but an H shape would have a less steep slope as it has a wider tuning. We took the derivative of each cumulative sum over frequency curve and found the maximum value for each curve. We also took the cumulative sum across attenuation.

### Excitatory bandwidth analysis

To quantify the spread of activity across frequency, we calculated the binary receptive field sum (BRFS) for all neurons (Bowen, Winkowski et al. 2020). We used the binarized matrices calculated above and found the sum of significant responses at every sound level. We chose to use BRFS instead of a traditional bandwidth calculation which only considers the main receptive field so we could account for multipeaked neurons.

### Inhibitory sideband analysis

We analyzed the tuning and inhibitory properties of neurons to understand how frequency responses and inhibition varied across groups. To calculate tuning width, we used the neuron’s tuning curve, which represents its response magnitude across frequencies. Significant response frequencies were identified using statistical criteria from two-tone stimuli data, and the tuning width was defined as the weighted average of octave differences between these frequencies and the neuron’s best frequency (BF). Weights were determined by the response amplitudes, and disconnected significant regions were managed using a merging threshold to ensure consistent calculations. In addition to tuning width, we measured inhibitory width to assess the range of frequencies around the BF where significant inhibition occurred. This was further divided into low- and high-frequency inhibitory sidebands, calculated as weighted averages of inhibitory amplitudes below and above the BF, respectively. The total inhibitory width pooled all significant inhibitory regions to provide an overall measure.

To quantify the strength of inhibition, we calculated the amplitude of inhibition by summing the magnitudes of inhibitory responses in the significant sidebands. To ensure comparisons across neurons with varying response magnitudes, we normalized this value by the peak response amplitude of the tuning curve, resulting in a relative amplitude of inhibition. To facilitate comparisons of tuning and inhibitory properties across neurons with different BFs, we aligned tuning curves and inhibitory sideband data relative to each neuron’s BF. For each neuron, we shifted the data arrays such that the BF was centered in the array, with sufficient range to include frequencies on both sides of the BF. This alignment and normalization enabled direct comparisons across neurons and groups.

### Statistics

All significance tests were conducted using built in MATLAB functions. 2 way ANOVAS were calculated using the anovan function. Student’s t tests between group means were conducted using the function ttest2 and multiple comparisons were corrected for using Bonferroni correction assuming all data is normally distributed.

## RESULTS

To characterize the changes of spectral processing that occur with aging, we imaged the primary auditory cortex (A1) of awake aging (12-24 months) and adult (11-24 weeks) mice, both male and female. We used F1 mice from a CBA/CaJ and Thy1GCaMP cross while playing sets of tones and two-tone complexes (Fig. 1a). We utilized a CBA background as these mice are known to retain peripheral hearing well into old age, allowing us to isolate aging effects to central areas (Willott, Parham et al. 1988, Willott, Parham et al. 1991, Bowen, Winkowski et al. 2020). We identified A1 using widefield imaging by playing pure tones, finding the areas that maximally responded to each tone, and comparing the resulting map to known tonotopic and anatomical markers typically used to locate A1 (Fig. 1b) (Liu, Whiteway et al. 2019). We imaged neurons in L2/3 of A1 under 2 photon imaging using a 512×512 window at 2x magnification (30Hz) (Fig. 1c) and extracted resulting fluorescent traces from neurons using suite2p (Pachitariu et al., 2016) and custom software.

**Figure 1.**
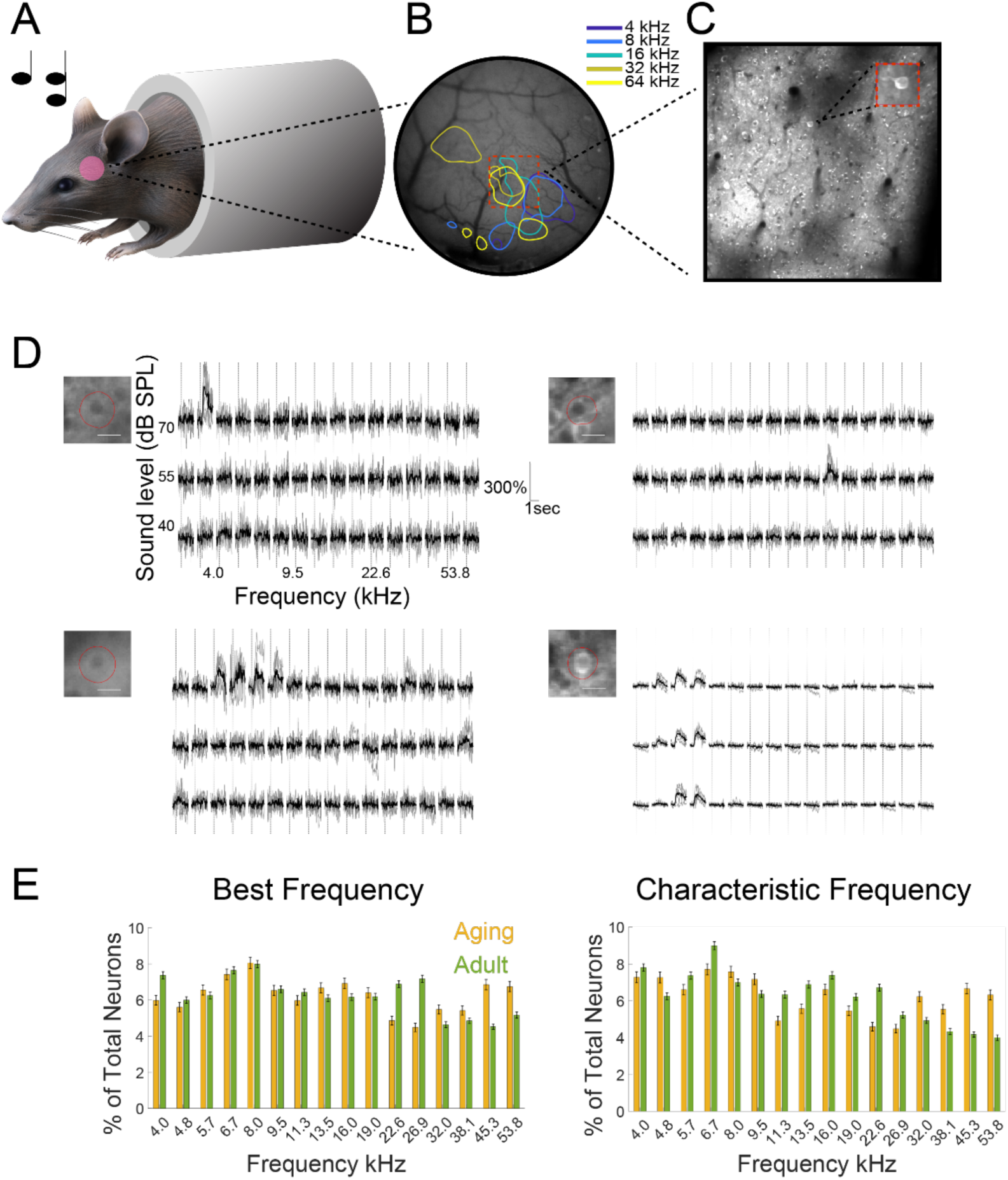
Experimental paradigm for in vivo imaging of auditory receptive fields. **A**. Example setup of imaging experiment. Cranial windows were surgically implanted over auditory cortex. Mice were headfixed inside of custom 3D printed tubes and passively listened to auditory stimuli during imaging. **B.** Widefield imaging combined with passive listening of pure tones was performed after cranial window surgery to identify tonotopic excitation patterns associated with known auditory fields. Colored patches show areas that significantly responded to different frequences played **C.** Typical 2x field of view in 2 photon imaging of primary auditory cortex. Typically, we were able to image 100-300 neurons simultaneously at 30 Hz. **D.** Example FRA responses to 16 different frequencies at 3 sound levels. For each FRA, the corresponding neuron is shown in an upper left panel and is outlined in red. 99% Confidence intervals are shown in light gray, the average response shown in black. Top row shows 2 neurons that respond sparsely to a specific sound level and frequency combination. Bottom row shows 2 neurons that have more complex receptive fields with responses spanning multiple frequencies and decibel levels. **E**. Best frequency and characteristic frequency distribution shown for aging vs adult animals. Best frequency and characteristic frequency were calculated for all neurons for both groups.

### A1 receptive fields in both aging and adult mice have variable shapes consistent with previous studies

Previous studies have identified a large degree in variability across receptive field shapes in multiple animal models including cats, rats, and mice (Abeles and Goldstein 1972, Sutter and Schreiner 1991, Sutter, Schreiner et al. 1999, Sutter 2000, Li, Liang et al. 2019, Liu and Kanold 2021). Additionally, a study in Fischer Brown-Norway rats (a model used in aging research) found that the distribution of frequency response area (FRA) types in layer 5 significantly changed with age (Turner, Hughes et al. 2005). To define our pure tone receptive fields, we presented a series of 16 pure tones (log spaced between 4 and 53.8 kHz) at 3 sound levels (40/55/70 dB) and plotted the average evoked fluorescence during the stimulus presentation as a function of frequency and sound level yielding an FRA. Only neurons that had at least one significant response at any frequency/decibel combination were included in this and further analyses. Significance was determined by using bootstrap analysis to find confidence intervals (CI) of the pre and post stimulus window and verifying that there was no overlap of the upper CI of the pre stimulus period with the lower CI of the post stimulus period. Here, FRAs are shown for 4 example neurons in figure 1d. We observed a large variety in sound-evoked responses across neurons, ranging from responses that cover multiple frequencies and/or decibel levels, to sparse responses that only cover one frequency/decibel combination.

### Best frequency and characteristic frequency distributions remain unchanged with age

Mice on a CBA/CaJ background have been shown to retain high frequency hearing in old age (Willott, Parham et al. 1988, Spongr, Flood et al. 1997, Bowen, Winkowski et al. 2020, Shilling-Scrivo, Mittelstadt et al. 2021). To validate this for the present study, we calculated the percentage of total neurons for each best and characteristic frequency. A neuron’s best frequency is the frequency that elicits the strongest response at the highest decibel level while the neuron’s characteristic frequency is the frequency it is most sensitive to at the lowest tested decibel level. We plotted the proportion of neurons across all tested frequencies as histograms for both cases and found no significant difference in the percentage of neurons responding to higher frequencies between the adult and aging groups (Fig. 1e). Therefore, the aging mice in our study did not exhibit marked changes in high frequency representation, which is consistent with previous studies (Shilling-Scrivo, Mittelstadt et al. 2021).

### Unsupervised Classification of FRAs reveals 5 FRA types in aging and adult mice

In adult animals, A1 neurons fall into distinct response classes based on their spectral response properties (Liu and Kanold 2021). We thus set out to investigate if neurons from aging animals also showed these response classes and if the distribution of neurons over these classes was altered with aging. In adult animals the receptive fields (e.g. the FRA) of layer 2/3 neurons show multiple characteristic shapes (Liu and Kanold 2021). Given that neurons have different best frequencies (BFs) each response shape can occur at varying parts of the stimulus space (e.g. at different best frequencies). To compare FRA shapes across frequency and across age we first aligned each cell’s FRA to its geometric center. While neurons were manually clustered in the study we were mimicking (Liu and Kanold 2021), we chose to use an unsupervised clustering method to remove any bias from our FRA classification. We used Principal component analysis (PCA) to reduce the dimensionality of the dataset and found that the top 6 components were sufficient to explain 95% of the variance, as marked by the green diamond (Fig2a). We then performed K-means clustering on the reduced-dimension data. When inspecting the sum of squares elbow plot, we found points 4 and 5 at the crux of the elbow (Fig 2b). The corresponding clusters and t-SNE plot are shown in Fig2c with the top 4 clusters in row 1 (corresponding to green diamond in Fig 2B) and the top 5 clusters in row 2 (corresponding to pink diamond in Fig 2B). Though both points were on the elbow, we chose to use 5 clusters for our subsequent data analysis, both to use a number that captured more variance in the dataset and hence, stricter criteria, and because 5 clusters sorted the FRA shapes into biologically meaningful and distinct categories similar to prior analysis (Liu and Kanold 2021). This suggests that neurons in aging animals retain a somewhat similar variety in tuning properties represented as diverse FRA shapes as previously reported.

**Figure 2.**
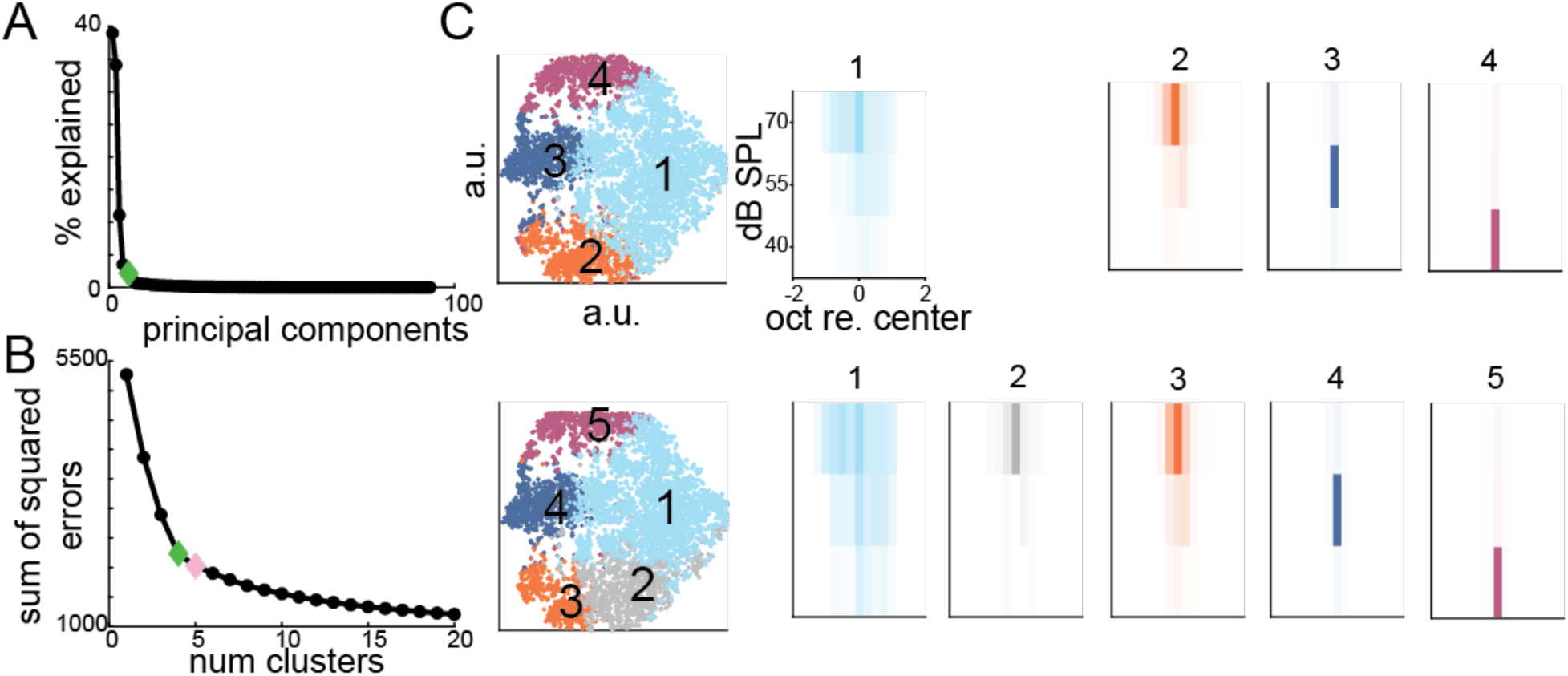
Unsupervised clustering of FRAs reveals 5 distinct cluster shapes. **A.** Elbow plot showing number of PCs vs variance explained in the dataset. The “elbow” of the curve occurred at 6 PCs (green diamond) which is when 95% of the variance is explained. The reduced dataset was then fed into the Kmeans clustering algorithm. **B.** Sum of squares elbow plot “elbow” occurred at 4 clusters (green diamond), however, 5 clusters (pink diamond) were chosen to capture more variance. **C.** T-SNE plot (top row) of 4 Kmeans clusters as suggested by the elbow plot in **B**. The bottom row shows the distribution of data and FRA types when dividing the data into 5 clusters. 5 clusters were ultimately chosen as they both display distinct FRA categories can account for more variance.

### PCA and K-means clustering reveal sex and age differences among FRA types

Several studies on aging have reported sex differences in both circuit makeup and functional response properties (Shilling-Scrivo, Mittelstadt et al. 2021, Shilling-Scrivo, Mittelstadt et al. 2022, Xu, Xue et al. 2025). To explore differences between age and sex, we split our clustered dataset into 4 groups of adult female, adult male, aging female, and aging male (Fig3A). For each sub figure, the t-SNE plot of the PCA reduced dataset is shown on the left followed by the top 5 clusters and their corresponding FRAs shown plotted to the right. We did not find any striking differences in the t-SNE plots across either sex or age but did notice distortions between the aging and adult FRA shapes particularly in the V, H, and I clusters. When plotting in t-SNE space, only local relationships among data are preserved; global relationships cannot be inferred. Hence, we can explore data similarity within clusters to explore changes across age and sex. We quantified these differences by computing a compactness score for each cluster where we found the Euclidian distance between each point and the center of each cluster and found the average of all distances per cluster. We found that, when comparing all animals for all FRA types, the compactness score was significantly higher for aging females within the S2 cluster and in aging females in the S3 cluster (Fig3B, student’s ttest adult vs aging with Bonferroni multiple comparison correction Adult/Aging V: *p=*0.03, H: *p=*0.09, I: *p=*0.15, S2: *p=*0.20, S3: *p=*0.11; Adult female/Aging female: V: *p=*5.58×10-3, H: *p=*0.19, I: *p=*0.26, S2: *p=*0.39, S3: *p=*0.03; Adult male/Aging male: V: *p=*0.33, H: *p=*0.14, I: *p=*0.25, S2: *p=*0.55, S3: *p=*0.45). This suggests that in aging animals, the neurons within S2 and S3 clusters become more heterogeneous, however, we cannot extrapolate from t-SNE *which* features are more diverse (i.e. bandwidth, shape, etc).

### Proportion of sparsely responding neurons to overall population increases in aging mice

In adult CBA/CaJxThy1-GCaMP6s mice, sparsely responding neurons make up approximately 70% of the FRA types found in L2/3 of auditory cortex while the other 30% are broadly tuned shapes (Liu and Kanold 2021). Aging increases the proportion of sparsely responding FRA types in L5 of the FSB rat auditory cortex and decreases the proportion of broadly tuned FRA types (Turner, Hughes et al. 2005). To investigate in detail how aging changes the FRA population makeup in our study, we calculated the proportion of neurons in each FRA category in the overall population for both sex and age categories (Fig 3C). We found a lower proportion of sparsely tuned neurons, especially S1 neurons, (roughly 28% vs. 50%) compared to prior work (Liu and Kanold 2021). We here clustered aging and adult neurons together, thus if the aging population are different enough, they will bias the clusters away from revealing a distinct S1 shape and instead these neurons may be grouped in the V, H, or I clusters. We did find a significant increase in the proportions of both S2 and S3 cluster types in aging animals (Student’s t test with Bonferroni multiple comparisons correction: p= Adult/Aging V:0.43 H:0.28, I:0.34 S2:3.41×10-4, S3:0.055; Adult female/Aging female: V:0.59, H:0.43, I:0.16, S2:0. 02, S3:0.01; Adult male/Aging male V:0.68, H:0.57, I:0.16, S2:1.6×10-3, S3:0.50246) which is consistent with findings in rat L5 (Turner, Hughes et al. 2005). Together these results suggest that aging increases the proportion of neurons with sparse FRAs, suggesting that either functional connections to non-sparse FRA neurons are lost with aging or that such neurons disappear.

**Figure 3.**
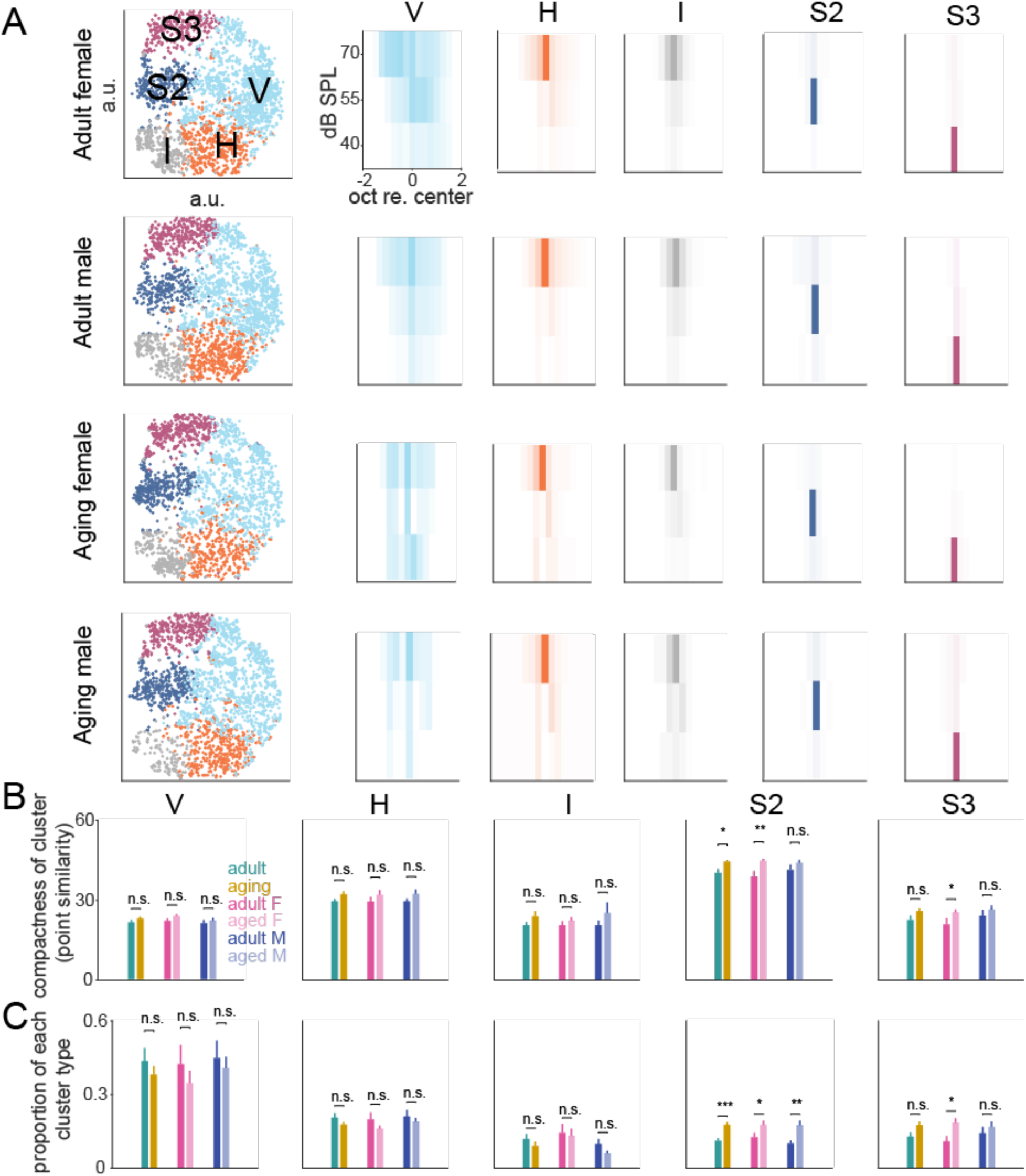
Aging reduces intra cluster similarity and alters proportion makeup in select cluster types **A.** Left column, t-SNE plot of PCA coefficients of center-aligned FRAs. Colors match the FRA cluster shapes in the 5 right columns found using Kmeans clustering. PCA, Kmeans, and t-SNE were applied to the entire dataset, then subgroups were separated out and plotted separately as adult female, adult male, aging female, and aging male. **B.** Compactness scores are plotted for each cluster subgroup. We calculated a compactness score as a proxy of point similarity by finding the Euclidean distance of each point from the center and then taking the mean of all scores. A higher compactness score means greater spread of points, hence, lower overall similarity of points to each other. Cluster types S2 and S3 were the only types to show significant differences. (Student’s t test with Bonferroni multiple comparisons correction: *p*= Adult/Aging V:0.03, H:0.09, I:0.15, S2:0.20, S3:0.11; Adult female/Aging female: V:5.58×10^-3^, H:0.19, I:0.26, S2:0.39, S3:0.03; Adult male/Aging male: V:0. 33, H:0.14, I:0. 25, S2:0. 55, S3:0.45). **C.** Proportion of neurons in each cluster per group. Both aging males and females showed an increase in S2 cluster types and aging females additionally showed an increase in S3 cluster type. (Student’s t test with Bonferroni multiple comparisons correction: *p*= Adult/Aging V:0.43 H:0.28, I:0.34 S2:3.41×10^-4^, S3:0.055; Adult female/Aging female: V:0.59, H:0.43, I:0.16, S2:0. 02, S3:0.01; Adult male/Aging male V:0.68, H:0.57, I:0.16, S2:1.6×10^-3^, S3:0.50246). * represents p<=0.05, ** represents p<=1×10^-2^, *** represents p<=1×10^-3^

### Aging animals display increased variability in bandwidth makeup over frequency and sound level when compared across neurons

The receptive fields of A1 neurons are formed by a complex interplay of both inhibitory and excitatory inputs (Tan, Zhang et al. 2004, Zhou, Liang et al. 2014, Meng, Winkowski et al. 2017, Moore, Weible et al. 2018, Liu and Kanold 2021). Studies in aging have found that both excitatory and inhibitory mechanisms degrade including reduced GAD levels, a reduction in inhibitory inputs, and overall changes in connectivity (Willott, Aitkin et al. 1993, Chaudhry, Reimer et al. 1998, Milbrandt, Holder et al. 2000, Shi, Argenta et al. 2004, Liao, Han et al. 2016, Kumar, Thinschmidt et al. 2019, Xu, Xue et al. 2025). Our clustering analysis suggested that there is a higher variability of sparsely tuned neurons from aging animals. We next sought to quantify and visualize the extent of the variability in responses in frequency or sound level across the entire receptive field from neurons in adult and aging animals. We first compared responses across animals (Fig 4B, D) and found significant differences between aging and adult in V and H type clusters in that the slope for aging animals was steeper than in adult – suggesting greater variability across frequency space in adult animals (Student’s t test between derivative distributions with Bonferroni multiple comparisons correction: p= Adult/Aging V:2.14×10^4^, H:9.3×10^-3^, I:0.24, S2:0.31, S3:0.41). No significant differences were observed across SPL space. However, when analyzing individual neurons, all cluster types showed steeper cumulative frequency curves in aging animals, suggesting narrower frequency response ranges. (Student’s t test with Bonferroni multiple comparisons correction: p= Adult/Aging V:0, H:0, I:0, S2:0, S3:0). In contrast, cumulative responses across SPL revealed greater representation of lower SPLs in aging neurons, even in clusters typically driven by high SPLs (Adult vs aging: 2-way ANOVA with post hoc ttest: p= V 70 SPL: 1.89×10^-230^, 55 SPL: 3.59×10^-188^, 40 SPL: 2.43×10^-134^; H 70 SPL: 5.82×10^-288^, 55 SPL: 1.57×10^-174^, 40 SPL: 2.66×10^-105^; I 70 SPL: 5.45×10^-180^, 55 SPL: 1.51×10^-95^, 40 SPL: 4.73×10^-45^; S2 70 SPL: 9.79×10^-165^, 55 SPL: 3.57×10^-185^, 40 SPL: 1.48×10^-52^; S3 70 SPL: 1.29×10^-146^, 55 SPL: 1.30×10^-174^, 40 SPL: 2.54×10^-205^). These findings suggest subtle age-related changes in frequency bandwidth and response distribution in all cell types.

**Figure 4.**
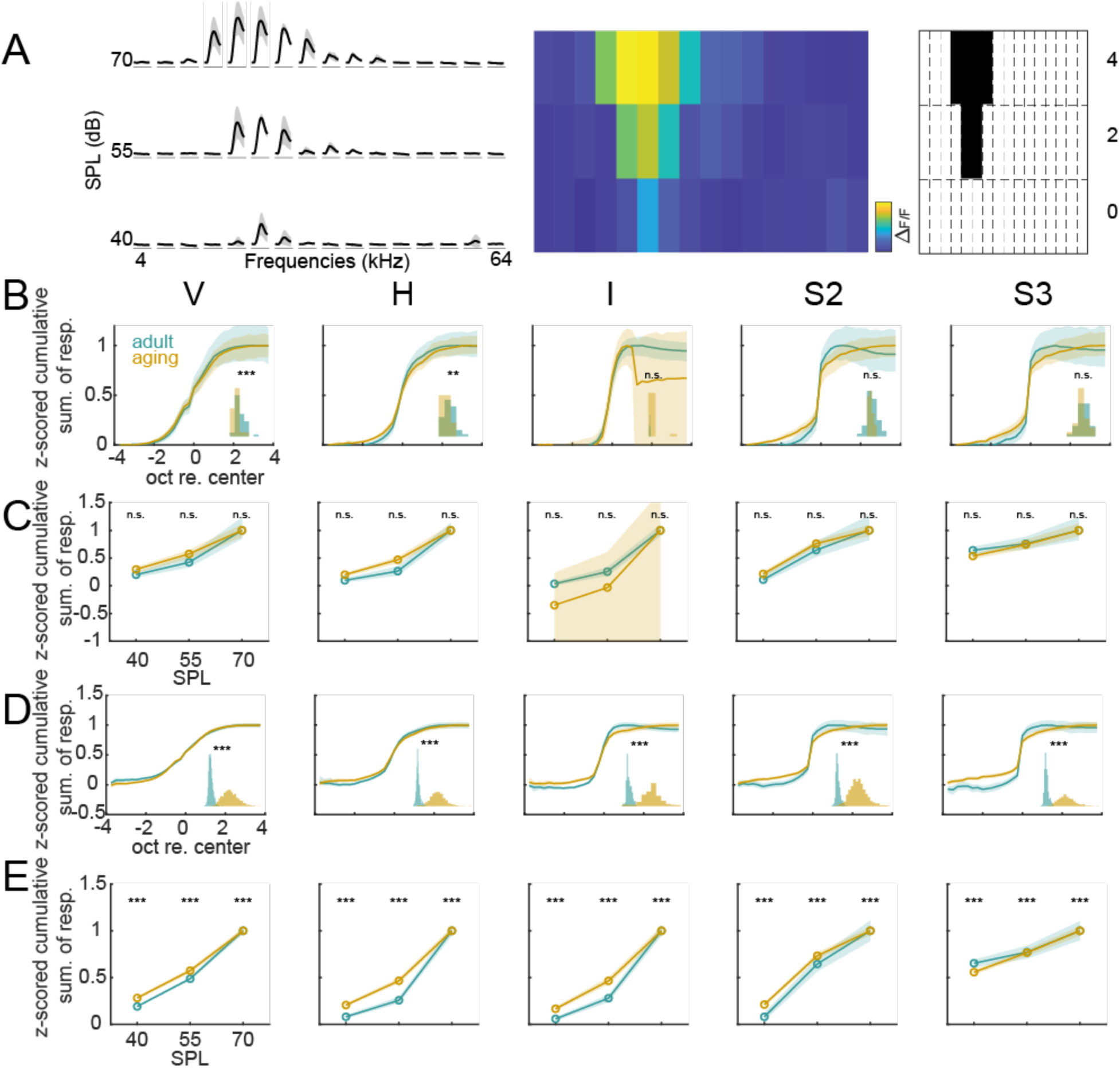
Aging alters excitatory responses across frequency and SPL **A.** Cartoon illustrating bandwidth calculation. Example FRA illustrated on left (10 point moving average for visual purposes, not included in bandwidth calculation). We first took the maximum value of each curve to create a 2D matrix of responses (middle plot). We then binarized the matrix, setting all responses above 30% of the max to 1 and all those below to 0. **B.** Cumulative responses over frequency are shown for adult and aging animals. Solid lines represent the z-scored responses of neurons in each FRA shape class and shaded regions represent 95% confidence intervals. We took the maximum derivative across neurons for each curve and plotted the distributions as a histogram for each FRA type in the lower right corner of each subplot. (Student’s t test: p= Adult/Aging V:2.14×10^4^, H:9.3×10^-3^, I:0.24, S2:0.31, S3:0.41) **C.** Cumulative responses over decibel level across animals are shown for adult and aging. Here, the matrix of binarized responses was collapsed across frequency to form a vertical vector. Then the responses were added up across frequencies, the z score was taken, and each additive step was plotted. Solid lines represent the z-scored responses of neurons in each FRA shape class and shaded regions represent 95% confidence intervals. (Adult vs aging: 2-way ANOVA with post hoc ttest: p= V 70 SPL: 1.89×10^-230^, 55 SPL: 3.59×10^-188^, 40 SPL: 2.43×10^-134^; H 70 SPL: 5.82×10^-288^, 55 SPL: 1.57×10^-174^, 40 SPL: 2.66×10^-105^; I 70 SPL: 5.45×10^-180^, 55 SPL: 1.51×10^-95^, 40 SPL: 4.73×10^-45^; S2 70 SPL: 9.79×10^-165^, 55 SPL: 3.57×10^-185^, 40 SPL: 1.48×10^-52^; S3 70 SPL: 1.29×10^-146^, 55 SPL: 1.30×10^-174^, 40 SPL: 2.54×10^-205^) **D.** Cumulative sum of responses over frequency across neurons. (Student’s t test: p= Adult/Aging V:0, H:0, I:0, S2:0, S3:0) **E.** Cumulative responses over decibel level across neurons are shown for adult and aging. (Adult vs aging: 2-way ANOVA with post hoc ttest p= all n.s.) * represents p<=0.05, ** represents p<=1×10^-2^, *** represents p<=1×10^-3^

### Excitatory bandwidth decreases across V, H, I shapes at the highest SPL level

Prior studies in aging mice found that in quiet, aging and adult mice had similar bandwidths but introducing a background of white noise, at low SNRs, decreased bandwidth in aging animals with similar results across sex (Shilling-Scrivo, 2021). However, these analyses were done across all neurons in the study grouped together and did not separate neurons by FRA shape. Identifying potential changes specific to FRA shapes can reveal information about functionally sub grouped neurons that are more prone to changes with aging. Here, we thus explored if there are changes specific to cluster type. For this analysis, we computed the binary receptive field sum (BRFS) of each FRA to include responses outside of the central response area (Bowen, Shilling-Scrivo et al. 2024). We then computed the average BRFS per age group, sex, and SPL stimulus level to compare excitatory bandwidth across groups. We found that at the highest SPL level, FRA types V, H, and I exhibited significant decreases in bandwidth but none at lower SPLs (Fig, 5a). We did not find significant differences for either of the sparsely tuned groups S2 or S3. These changes were present in both sexes (Fig. 5b, c) (Adult vs aging: Student’s t test with Bonferroni multiple comparisons correction: p= V 70 SPL: 7.5×10^-5^, 55 SPL: 0.07, 40 SPL: 0.34; H 70 SPL: 7.7×10^-3^, 55 SPL: 0.023, 40 SPL: 0.04; I 70 SPL: 1.55×10^-6^, 55 SPL: 0.09, 40 SPL: 0.12; S2 70 SPL: 0.12, 55 SPL: 0.87, 40 SPL: 0.21; S3 70 SPL: 0.99, 55 SPL: 0.14, 40 SPL: 0.81, Female adult vs female aging: Student’s t test with Bonferroni multiple comparisons correction: p= V 70 SPL: 7.6×10^-3^, 55 SPL: 0.06, 40 SPL: 0.42; H 70 SPL: 0.13, 55 SPL: 0.79, 40 SPL: 0.23; I 70 SPL: 5.1×10^-5^, 55 SPL: 0.06, 40 SPL: 0.39; S2 70 SPL: 0.27, 55 SPL: 0.13, 40 SPL: 0.05; S3 70 SPL: 0.92, 55 SPL: 0.52, 40 SPL: 0.35. Male adult vs male aging: Student’s t test with Bonferroni multiple comparisons correction: p= V 70 SPL: 4.0×10^-3^, 55 SPL: 0.63, 40 SPL: 0.58; H 70 SPL: 0.02, 55 SPL: 0.07, 40 SPL: 0.11; I 70 SPL: 7.7×10^-3^, 55 SPL: 0.57, 40 SPL: 0.21; S2 70 SPL: 0.3, 55 SPL: 0.35, 40 SPL: 0.74; S3 70 SPL: 0.9, 55 SPL: 0.14, 40 SPL: 0.41). These results suggest that while aging has an effect on excitatory bandwidth, it disproportionately impairs broadly tuned neurons and does not strongly affect sparsely tuned neurons likely because these neurons receive different inputs.

**Figure 5.**
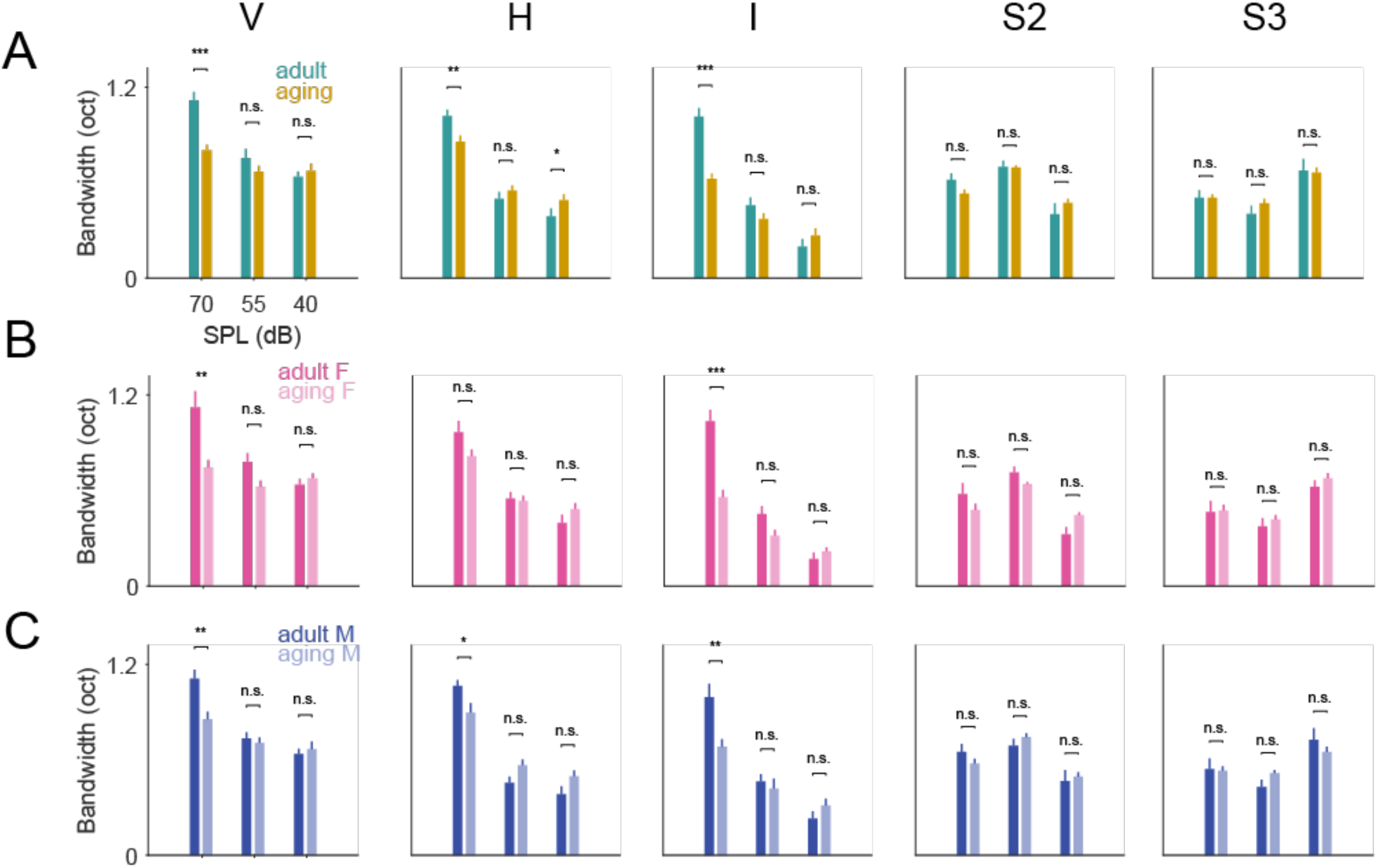
Excitatory bandwidth decreases with age **A.** Bandwidth was calculated by taking the binary receptive field sum. The leftmost plot of A shows the count of responses per frequency/decibel level. The count per decibel level was taken and averaged across neurons per animal. Results are shown for adult versus aging. (Adult vs aging: Student’s t test with Bonferroni multiple comparisons correction: p= V 70 SPL: 7.5×10-5, 55 SPL: 0.07, 40 SPL: 0.34; H 70 SPL: 7.7×10-3, 55 SPL: 0.023, 40 SPL: 0.04; I 70 SPL: 1.55×10-6, 55 SPL: 0.09, 40 SPL: 0.12; S2 70 SPL: 0.12, 55 SPL: 0.87, 40 SPL: 0.21; S3 70 SPL: 0.99, 55 SPL: 0.14, 40 SPL: 0.81 **B.** As in D, bandwidths are shown per cluster group for adult versus aging females. (Female adult vs female aging: Student’s t test with Bonferroni multiple comparisons correction: p= V 70 SPL: 7.6×10-3, 55 SPL: 0.06, 40 SPL: 0.42; H 70 SPL: 0.13, 55 SPL: 0.79, 40 SPL: 0.23; I 70 SPL: 5.1×10-5, 55 SPL: 0.06, 40 SPL: 0.39; S2 70 SPL: 0.27, 55 SPL: 0.13, 40 SPL: 0.05; S3 70 SPL: 0.92, 55 SPL: 0.52, 40 SPL: 0.35) **C**. Bandwidths are shown per cluster group for adult versus aging males. (Male adult vs male aging: Student’s t test with Bonferroni multiple comparisons correction: p= V 70 SPL: 4.0×10-3, 55 SPL: 0.63, 40 SPL: 0.58; H 70 SPL: 0.02, 55 SPL: 0.07, 40 SPL: 0.11; I 70 SPL: 7.7×10-3, 55 SPL: 0.57, 40 SPL: 0.21; S2 70 SPL: 0.3, 55 SPL: 0.35, 40 SPL: 0.74; S3 70 SPL: 0.9, 55 SPL: 0.14, 40 SPL: 0.41). * represents p<=0.05, ** represents p<=1×10^-2^, *** represents p<=1×10^-3^

### Absolute and relative inhibition weaken in aging animals

Inhibitory circuits shape the structure of receptive fields (Seybold, Stanco et al. 2012, Li, Ji et al. 2014, Seybold, Phillips et al. 2015, Kato, Asinof et al. 2017, Natan, Rao et al. 2017, Moore, Weible et al. 2018, Lakunina, Nardoci et al. 2020, Nocon, Gritton et al. 2023) and aging causes a decrease in suppressed responses in A1 suggesting decreased inhibition (Shilling-Scrivo, Mittelstadt et al. 2021). In order to explore the effect of inhibition on each class of FRA in more detail, and to reveal changes in inhibition in aging animals, we used two tone stimuli (Sachs and Kiang 1968, Abeles and Goldstein 1972, Nelken, Prut et al. 1994). Classically, in two-tone stimuli a tone at the neurons’ BF is played together with a second tone that can alter the cell’s response – this can be inhibitory or facilitative. Because we image 100s of neurons simultaneously, we cannot individually adjust our stimuli for every single neuron. Thus, our stimulus set consisted of pairs of the same 16 tones we used for our single tone experiment for a total of 120 tone pairs. Both pure tones were played at the same sound level, resulting in a combined 63 dB sound level. Two tone and single tone responses are shown for an example cell in Figure 6A (left panel) where the cell’s best frequency or BF is around 7 kHz. This example demonstrates that the two-tone paradigm can identify inhibitory sidebands in the auditory cortex.

**Figure 6.**
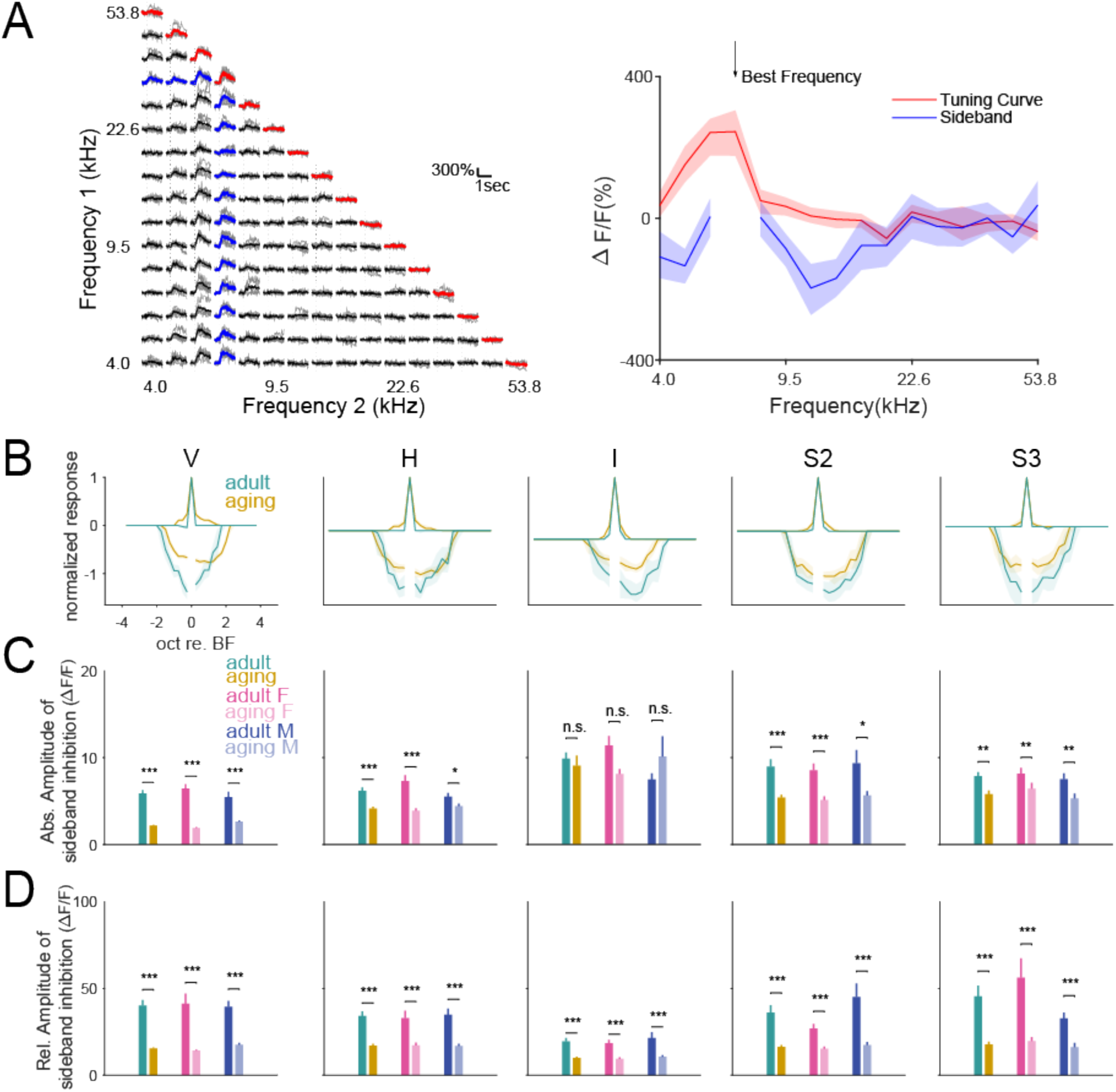
Inhibitory sidebands weaken in aging animals. **A**. Two tone combinations were presented to animals to assess sideband contribution to best frequency responses. Matrix shown of single tone responses (diagonal line in red) and two tone responses (all other responses in black or blue). Sum of responses are shown to the right. Shaded regions show confidence intervals **B.** Pooled responses across cluster groups. Traces above 0 indicate tuning curves, traces below 0 indicate inhibitory sidebands. All tuning curves were normalized to their maximum response. Inhibitory sidebands were normalized to the max of their respective tuning curves where no change in response would be a value of 0 and any inhibitory response would be negative. Shaded regions show confidence intervals. **C.** Absolute amplitude of sideband inhibition was measured by taking the absolute value of sideband dff and taking the average of all significant responses (Student’s t test with Bonferroni multiple comparisons correction: *p=* Adult/Aging V: 1.1×10^-31^, H: 1.8×10^-7^, I: 0.67, S2: 1.7×10^-4^, S3: 1.1×10^-3^; Adult female/Aging female: V: 6.6×10^-25^, H: 2.8×10^-6^, I: 0.05, S2: 3.2×10^-3^, S3: 9.3×10^-3^; Adult male/Aging male: V: 1.0×10^-6^, H: 0.03, I: 0.51, S2: 0.03, S3: 9.3×10^-3^). **D.** Relative amplitude of sideband inhibition consisted of normalizing the inhibitory response to the maximum tuning curve response. (Student’s t test with Bonferroni multiple comparisons correction: *p=* Adult/Aging V: 6.2×10^-24^, H: 1.2×10^-11^, I: 6.0×10^-10^, S2: 9.6×10^-6^, S3: 8.3×10^-5^; Adult female/Aging female: V: 1.0×10^-17^, H: 2.8×10^-8^, I: 1.3×10^-7^, S2: 2.6×10^-6^, S3: 3.1×10^-5^; Adult male/Aging male: V: 1.3×10^-10^, H: 4.0×10^-7^, I: 4.4×10^-5^, S2: 7.6×10^-4^, S3: 1.2×10^-4^). * represents p<=0.05, ** represents p<=1×10^-2^, *** represents p<=1×10^-3^

To understand the effect aging has on inhibitory activity, we compared the sideband activity across FRA shape. As in Fig 6A, for each cell, we normalized the tuning curve and the inhibitory sideband by the maximum value of the tuning curve. We then centered all curves and sidebands at the neuron’s best frequency (Fig 6B). To calculate the amplitude of sideband inhibition, we took the absolute value of every significant two-tone response and then summed all responses to find the absolute amplitude (Fig 6C). To calculate relative amplitude, we divided this number by the maximum tuning curve response to normalize the response to the tuning curve response. Thus, our measure of sideband amplitude is a combination of both sideband width and response amplitude. We found that absolute amplitude of inhibition decreased in all cluster types with age except for the I type (Student’s t test with Bonferroni multiple comparisons correction: *p=* Adult/Aging V: 1.1×10^-31^, H: 1.8×10^-7^, I: 0.67, S2: 1.7×10^-4^, S3: 1.1×10^-3^; Adult female/Aging female: V: 6.6×10^-25^, H: 2.8×10^-6^, I: 0.05, S2: 3.2×10^-3^, S3: 9.3×10^-3^; Adult male/Aging male: V: 1.0×10^-6^, H: 0.03, I: 0.51, S2: 0.03, S3: 9.3×10^-3^). We also found that the sideband inhibition, relative to that cell’s tuning curve decreased across both age and sex for all FRA types. (Student’s t test with Bonferroni multiple comparisons correction: *p=* Adult/Aging V: 6.2×10^-24^, H: 1.2×10^-11^, I: 6.0×10^-10^, S2: 9.6×10^-6^, S3: 8.3×10^-5^; Adult female/Aging female: V: 1.0×10^-17^, H: 2.8×10^-8^, I: 1.3×10^-7^, S2: 2.6×10^-6^, S3: 3.1×10^-5^; Adult male/Aging male: V: 1.3×10^-10^, H: 4.0×10^-7^, I: 4.4×10^-5^, S2: 7.6×10^-4^, S3: 1.2×10^-4^). This suggests that aging reduces the strength of inhibitory sidebands globally in all clustered FRA types.

### Aging has greater widespread effect on inhibitory amplitude over bandwidth

To investigate changes in inhibition in detail, we next explored which aspects of inhibition are most affected – response amplitude or inhibitory sideband width. Since inhibitory sidebands can have variable shapes, we calculated the inhibitory sidebands width using the BRFS. Maximum inhibitory amplitude was found by taking the absolute value of the greatest response change. We found that only 2 cluster types, S3 and I, showed a significant decrease in inhibitory BRFS (Fig.7a), but that amplitude was decreased in almost all cluster types (Fig. 7b) (Inhibitory sideband bandwidth: Student’s t test with Bonferroni multiple comparisons correction: p= Adult/Aging V: 0.7, H: 0.94, I: 3.2×10-4, S2: 0.05, S3: 3.75×10-5; Adult female/Aging female: V: 0.19, H: .48, I: 0.01, S2: 0.07, S3: 6.2×10-4; Adult male/Aging male: V: 0.43, H: 0.72, I: 0.01, S2: 0.76, S3: 3.0×10-4). (Maximum inhibitory sideband value: Student’s t test with Bonferroni multiple comparisons correction: p= Adult/Aging V: 2.8×10^-36^, H: 1.1×10^-9^, I: 0,87, S2: 4.1×10^-4^, S3: 2.3×10^-2^; Adult female/Aging female: V: 1.0×10^-9^, H: .1.7×10^12^, I: 0.53, S2: 1.2×10^-4^, S3: 0.09; Adult male/Aging male: V: 4.2×10^-7^, H: 0.01, I: 0.32, S2: 0.04, S3: 0.09). These results demonstrate that aging has a significantly larger effect on the maximum amplitude of inhibition, but not on the bandwidth of inhibitory frequency inputs. This suggests that even though aging animals retain their sideband inhibition, they experience less robust sideband inhibition thus possibly less sharp tuning to their preferred frequency.

**Figure 7.**
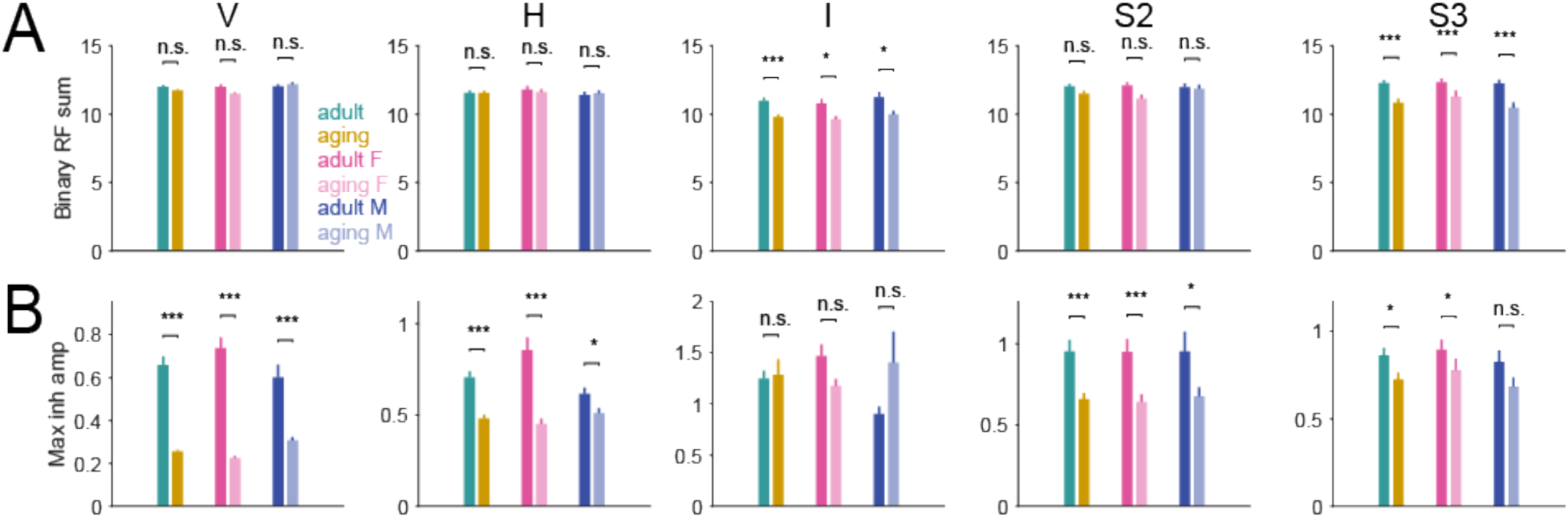
Aging has greater widespread effect on inhibitory amplitude over bandwidth **A.** Inhibitory sideband bandwidth was calculated by taking the binary receptive field sum or the sum of the number of significant inhibitory responses across the receptive field. (Inhibitory sideband bandwidth: Student’s t test with Bonferroni multiple comparisons correction: p= Adult/Aging V: 0.7, H: 0.94, I: 3.2×10-4, S2: 0.05, S3: 3.75×10-5; Adult female/Aging female: V: 0.19, H: .48, I: 0.01, S2: 0.07, S3: 6.2×10-4; Adult male/Aging male: V: 0.43, H: 0.72, I: 0.01, S2: 0.76, S3: 3.0×10-4) **B.** Maximum absolute value of inhibitory sideband responses(Maximum inhibitory sideband value: Student’s t test with Bonferroni multiple comparisons correction: p= Adult/Aging V: 2.8×10-36, H: 1.1×10-9, I: 0,87, S2: 4.1×10-4, S3: 2.3×10-2; Adult female/Aging female: V: 1.0×10-9, H: .1.7×1012, I: 0.53, S2: 1.2×10-4, S3: 0.09; Adult male/Aging male: V: 4.2×10-7, H: 0.01, I: 0.32, S2: 0.04, S3: 0.09). * represents p<=0.05, ** represents p<=1×10^-2^, *** represents p<=1×10^-3^

## DISCUSSION

We investigated how aging affects excitatory and inhibitory properties of receptive fields in primary auditory cortex (A1), focusing on the changes in receptive field shape, inhibitory sidebands, and variability. Use of CBA mice, which retain peripheral hearing in old age (Willott, Parham et al. 1988, Bowen, Winkowski et al. 2020) allowed us to isolate central mechanisms. Our results demonstrate that A1 undergoes profound changes in both the diversity of receptive field types and strength of inhibition during aging. We find that the numbers of distinct receptive field classes are reduced, the diversity of receptive fields in each class increases, excitatory bandwidth decreases, and that inhibitory sidebands weaken.

Unsupervised clustering of frequency response areas (FRAs) using PCA and K-means revealed five distinct response types in A1 layer 2/3 (L2/3), consistent with previous reports in mice and cats (Abeles and Goldstein 1972, Sutter and Schreiner 1991, Liu and Kanold 2021). These included sparsely tuned (S2, S3) and broadly tuned (V, H, I) classes. Notably, we did not observe the S1 subtype, which previously accounted for nearly 40% of adult neurons (Liu and Kanold 2021). S1 neurons are sparsely tuned and respond preferentially at higher SPLs. Their absence may reflect age-related degradation of receptive field boundaries, where formerly distinct S1 responses become more spectrally diffuse and merge with other categories. Because clustering was performed on pooled adult and aging data, degraded S1 responses may have shifted cluster boundaries, affecting both groups. Increased spectral overlap and loss of tuning specificity are hallmarks of aging auditory neurons (Willott, Parham et al. 1988, Krukowski and Miller 2001, Bao, Chang et al. 2004, Shilling-Scrivo, Mittelstadt et al. 2021). We also observed a shift in FRA type distribution with age. While prior studies found ∼70% of L2/3 neurons in adult mice to be sparsely tuned (40% S1, 30% S2/S3), our aging cohort lacked S1 responses but showed increased proportions of S2 and S3. This redistribution within sparse types, alongside stable proportions of V, H, and I neurons, suggests that age-related degradation disproportionately affects certain subtypes without uniformly impacting all FRA categories. These findings align with observations in aging rats, where sparse tuning becomes more common and broad FRA types decline (Turner, Hughes et al. 2005). The absence of S1 in our data may thus reflect a merging of degraded responses into broader or neighboring categories rather than a complete loss of sparsely tuned activity.

Increased within cluster variability was supported by t-SNE analysis, which revealed that S2 and S3 clusters were more dispersed in aging females, suggesting greater intra-cluster variability. Such variability may arise from differential susceptibility of cell types to age-related circuit changes. The mature auditory cortex relies on a delicate balance between excitatory and inhibitory inputs (Tan, Zhang et al. 2004, Zhou, Liang et al. 2014, Meng, Winkowski et al. 2017, Moore, Weible et al. 2018, Liu and Kanold 2021), and disruption of this balance—such as by silencing interneurons—triggers compensatory shifts in network dynamics (Moore, Weible et al. 2018). However, this homeostasis appears impaired with age, as in aging CBA mice, slice recordings show a shift toward excitation and reduced feedforward input from Layer 4 to L2/3, particularly in females (Xu, Xue et al. 2025).

Receptive field cluster degradation may selectively manifest in certain neuronal subtypes. In rat A1, distinct classes of Layer 5 pyramidal neurons differ in size, tuning sharpness, and susceptibility to GABAergic inhibition (Hefti and Smith 2000, Hefti and Smith 2003), with smaller, more inhibition-sensitive neurons displaying sparse tuning and greater vulnerability to aging (Turner, Hughes et al. 2005, Turner, Hughes et al. 2005). Analogously, mouse L2/3 neurons receive variable input from different layers depending on subtype (Meng, Winkowski et al. 2017), suggesting that FRA types like S2 and S3 may reflect circuit-specific vulnerabilities. The disappearance of S1 and increased variability in S2/S3 neurons may represent the collapse of a subtype once stabilized by precise excitatory-inhibitory balance.

Our derivative-based analyses of the FRA curves indicated greater diversity in response slopes across the frequency axis among aging neurons. Steeper derivatives suggested that fewer frequencies elicited strong responses, while a broader spread in derivative values point to increased variability across neurons. This analysis revealed that aging FRAs displayed much steeper slopes suggesting that there were fewer responses across frequency space for any FRA subtype but each had a much wider variety of maximum derivative values, suggesting that neuronal responses in aging mice across frequency space were more variable than FRAs in adult mice. This increased variance is consistent with prior observations in Layer 5 neurons in aging rats (Turner, Hughes et al. 2005) where neurons in aging animals were found to have less response reliability. When comparing across SPL levels, we found that aging animals almost always had greater responses at lower SPL levels than adult animals though this effect was only significant when comparing across neurons.

We examined inhibitory function using two-tone suppression, a method that provides frequency-specific assessment of lateral inhibition. Our results revealed age-related declines in both absolute and relative inhibition, with most FRA types affected. These findings align with reports that lateral inhibition is crucial for frequency selectivity and is impaired in aging (Willott, Aitkin et al. 1993, Kumar, Thinschmidt et al. 2019). The observed decline in inhibitory strength was present in both sexes, differing from earlier work that reported sex-specific effects in suppressive responses in aging (Shilling-Scrivo, Mittelstadt et al. 2021). Methodological differences may account for this discrepancy. Whereas previous studies used background noise to probe inhibitory responses, our two-tone design enabled us to directly measure sideband effects from individual frequencies. This approach may explain our detection of global inhibition loss across sexes, particularly in sidebands, though notably, while they observed sex-dependent effects when comparing inhibitory bandwidth, they did report reduced suppressive responses across sexes when clustering neural trace shapes—more consistent with our findings.

The main interneuron subtypes in the auditory cortex are Somatostatin (SST) neurons (Kato, Asinof et al. 2017, Lakunina, Nardoci et al. 2020) and Parvalbumin (PV) neurons (Li, Ji et al. 2014) which are known to decline with age (Ouellet and de Villers-Sidani 2014). PV inputs are especially dense in Layer 4, which sends strong feedforward input to L2/3. If PV-mediated inhibition is reduced in Layer 4, neurons in L2/3 may receive less refined input, leading to more variable or less selective tuning. On the circuit level, aging leads to reduced intra- and interlaminar excitatory inputs and fewer inhibitory contacts (Xu, Xue et al. 2025), consistent to increased variability and less consistent suppression of irrelevant stimuli.

We observed a reduction in excitatory bandwidth in aging animals, but only in broadly tuned V, H, and I types. This finding is consistent with prior results (Shilling-Scrivo, Mittelstadt et al. 2021), where bandwidth was shown to decrease when tones were presented against a background of noise in aging mice. Inhibition sharpens tuning curves in auditory cortex (Li, Ji et al. 2014), thus decreased inhibition would be expected to cause wider bandwidth. However, chronic decreases of cortical inhibition have been shown to lead to decreased excitatory bandwidth (Seybold, Stanco et al. 2012). Since aging is a slow process, our results are consistent with these studies and suggest that the excitatory bandwidth narrows over time which may be due to homeostatic pruning or reorganization of inputs, particularly those supporting broadly tuned neurons. While it is possible that a much finer frequency spacing could reveal bandwidth changes in sparse neurons, we speculate that the cell type specific changes we observe reflect differences in circuit makeup which may be differently affected by aging.

While earlier studies have emphasized temporal coding deficits in aging mice under noisy or active conditions (Shilling-Scrivo, Mittelstadt et al. 2021, Shilling-Scrivo, Mittelstadt et al. 2022), our findings focus on spectral encoding in passive conditions. The observed decrease in distinct clusters and increased variability within sparse types suggest that spectral coding is also compromised. The loss of broadly tuned neurons and reduction in bandwidth may hinder the ability to track complex or dynamic stimuli—essential features for speech-in-noise perception. A broader bandwidth may offer more flexibility for integrating natural sounds which are spectrally rich, and a decrease in this bandwidth may reduce perceptual robustness in older adults (Dubno, Dirks et al. 1984, Fitzgibbons and Gordon-Salant 1998, Gordon-Salant and Fitzgibbons 1999, Lister, Besing et al. 2002, Lee 2015). This interpretation is supported by literature on spectrotemporal complexity in natural sounds and the importance of distributed tuning for adaptive auditory processing (Smolders, Aertsen et al. 1979, Nelken, Rotman et al. 1999, Bowen, Shilling-Scrivo et al. 2024).

Taken together, our findings suggest that the aging auditory cortex undergoes a multifaceted transformation affecting both excitatory and inhibitory components of sound processing. This includes an increase in tuning variability, a shift in FRA type distributions, a decrease in excitatory bandwidth, and a degradation of inhibitory function. These changes likely have substantial implications for how spectrally complex auditory information is encoded and processed at higher levels.

## Acknowledgements

Supported by NIH, P01AG55365 (POK), R21MH116450 (POK), R01DC17785 (POK), RF1AG078378 (POK).

